# ChAlPred: A Web Server for Prediction of Allergenicity of Chemical Compounds

**DOI:** 10.1101/2021.05.21.445101

**Authors:** Neelam Sharma, Sumeet Patiyal, Anjali Dhall, Naorem Leimarembi Devi, Gajendra P. S. Raghava

## Abstract

Allergy is the abrupt reaction of the immune system that may occur after the exposure with allergens like protein/peptide or chemical allergens. In past number of methods of have been developed for classifying the protein/peptide based allergen. To the best of our knowledge, there is no method to classify the allergenicity of chemical compound. Here, we have proposed a method named “ChAlPred”, which can be used to fill the gap for predicting the chemical compound that might cause allergy. In this study, we have obtained the dataset of 403 allergen and 1074 non-allergen chemical compounds and used 2D, 3D and FP descriptors to train, test and validate our prediction models. The fingerprint analysis of the dataset indicates that PubChemFP129 and GraphFP1014 are more frequent in the allergenic chemical compounds, whereas KRFP890 is highly present in non-allergenic chemical compounds. Our XGB based model achieved the AUC of 0.89 on validation dataset using 2D descriptors. RF based model has outperformed other classifiers using 3D descriptors (AUC = 0.85), FP descriptors (AUC = 0.92), combined descriptors (AUC = 0.93), and hybrid model (AUC = 0.92) on validation dataset. In addition, we have also reported some FDA-approved drugs like Cefuroxime, Spironolactone, and Tioconazole which can cause the allergic symptoms. A user user-friendly web server named “ChAlPred” has been developed to predict the chemical allergens. It can be easily accessed at https://webs.iiitd.edu.in/raghava/chalpred/.

## Introduction

Allergy is an inappropriate reaction of the immune response when it misidentifies a harmless foreign substance as a threat.^**1**^ These foreign substances are known as allergens, which could trigger several allergic reactions and lead to various allergic diseases. Different types of aeroallergens (e.g., pollens, spores, dust mites), food allergens (e.g., eggs, peanuts, tree nuts, genetically modified foods), and chemical allergens in personal care products (e.g., fragrances in the skin and hair care products, dyes, creams)^**1, 2, 3**^ can lead to allergic symptoms such as allergic asthma, rhinitis, skin reactions and anaphylaxis.^**1, 4**^ Anaphylactic shock involves a series of allergic reactions from mild symptoms like itchy skin, rashes, facial swelling, irritation of the eyes leading to watery eyes and nose to severe symptoms like shortness of breath, lack of consciousness, weak pulse, nausea, vomiting, which can even lead to death if untreated.^**5, 6**^ Certain studies show that allergic diseases are much more prevalent in developed countries than in developing countries.^**7, 8, 9, 10**^ In the last few years, the inflation in the occurrence of allergic diseases has increased the expenses of the treatment and also negatively influenced the status of life of a huge population.^**11**^

There is a wide variety of molecules that can pose a threat as allergens including biological molecules like proteins and peptides or some chemical compounds.^**1**^ In the past, several methods have been proposed to predict protein allergens from genetically engineered foods, vaccines and therapeutics. All the available methods are based on protein/peptide allergens, such as the recently developed method AlgPred 2.0.^**1, 12**^ Many other methods such as AllerTool^**13**^, AllerHunter^**14**^, AllerTOP^**15**^, AllerTOPv2^**16**^, PREAL^**17**^, AllergenFP^**18**^, AllerCatPro^**2**^ are heavily used by the scientific community. These methods are especially used in clinical researcher for designing proteins with desired allergenicity. In contrast, there is no method for predicting allergenic potential of the chemicals. Despite the fact, day-to-day life, the human body is exposed to innumerable chemical substances, such as makeup, soaps, perfumes, lotions, hair dyes, preservatives in food, metals in the jewellery.^**19**^ Many of these chemical products are known to provoke allergic reactions, causing skin sensitization in some people, which results in skin or contact dermatitis, and some may cause the sensitization of the respiratory tract leading to occupational asthma, which could be lethal.^**4, 5**^

Thus it is important to understand allergenicity of chemicals for developing methods for predicting allergic chemicals. Broadly, allergic reaction caused by small chemical compounds is developed in two phases; sensitization and elicitation. The first phase is initiated when a sensitive individual is exposed to a chemical allergen in sufficient amount, and via a proper route, then it will lead to immunological priming. In the context of allergy, immunological priming is called as sensitization or induction, which means that the mast cells and basophils are loaded with IgE antibodies against the chemical allergen.^**1, 20, 21**^ In the second phase, the re-exposure to the same chemical compound at the same or different site will provoke an accelerated and more aggressive secondary immune response. This secondary immune response is called as elicitation, which results in an allergic reaction. The already sensitized mast cells and basophils result in releasing cytoplasmic granules, and inflammatory molecules, such as, leukotriene, prostaglandins, histamine etc., leading to a mild allergic reaction to sudden death from anaphylactic shock.^**1, 20, 21**^ The mechanism of allergy caused by chemical allergens is depicted in Figure 1.

**Figure1:**
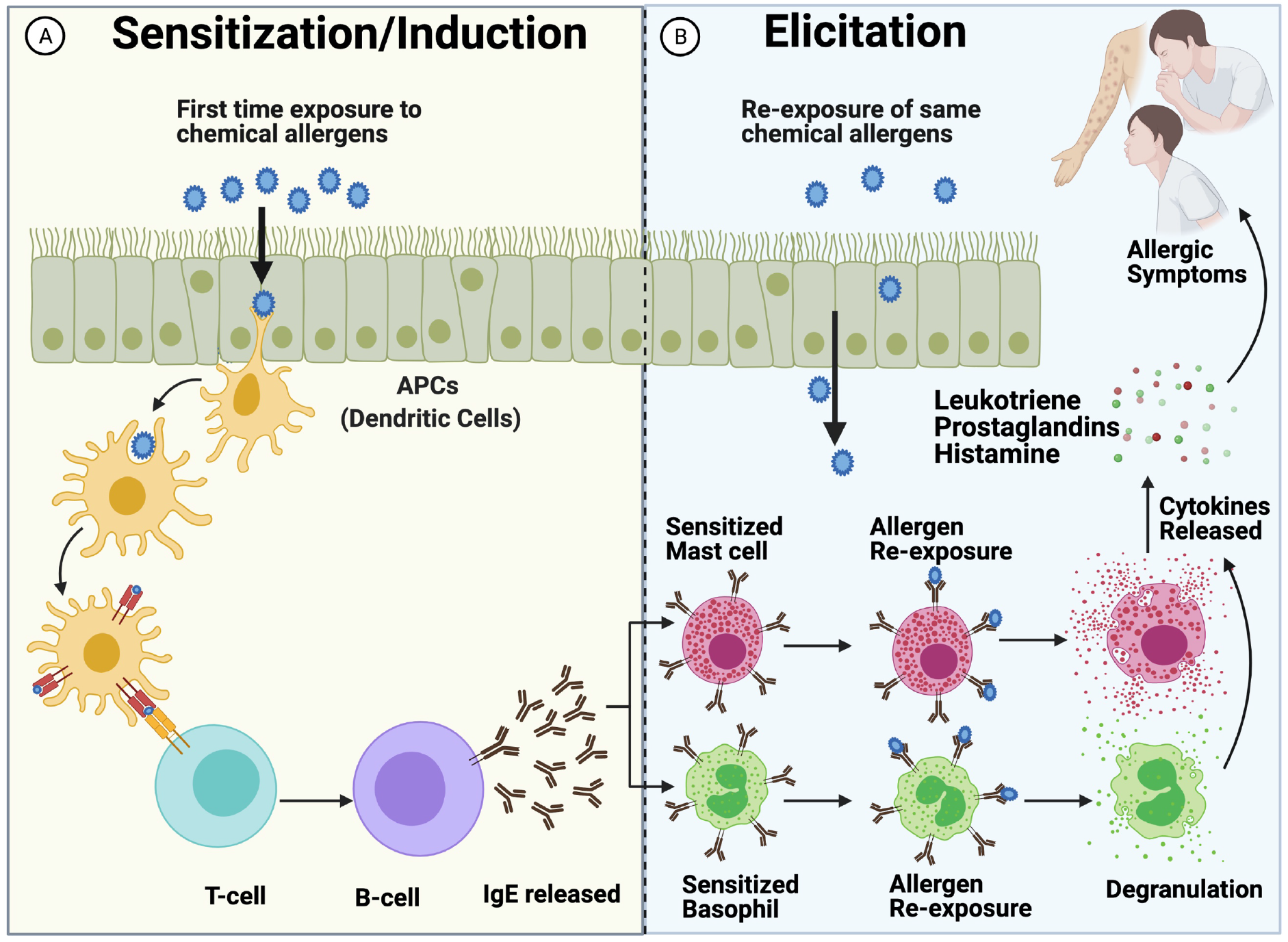
The mechanism of the allergy caused by chemical allergens.

In this study, first time systematic attempt had been made to develop in silico models for predicting allergic potential of chemicals. We obtained experimentally validated chemical-based allergen and non-allergens from well establish database IEDB. These chemicals were analyzed to understand chemical groups or finger prints responsible for causing allergenicity. In order to derive rules and to understand relationship between allergenicity and structure of chemicals, we compute wide range of descriptors using PaDEL software.^**22**^ These descriptors can be divided broadly in three categories; 2D descriptors, 3D descriptors and Fingerprints. Finally, we developed machine learning based models for predicting allergenicity of chemical using different type of descriptors. Our best models have been integrated into the webserver; it allows user to predict allergenicity of chemicals as well as allow to generate analogs of desired allergenicity https://webs.iiitd.edu.in/raghava/chalpred/.

## Materials and Methods

### Dataset Collection and Descriptors generation

In this study, we have collected allergenic and non-allergenic chemical compounds from the Immune Epitope Database (IEDB)^**23**^ and the structure for the same compounds were downloaded from Chemical Entities of Biological Interest (ChEBI) database.^**24**^ We obtained a total of 519 chemical compounds with allergenic properties from IEDB. On the other hand, we have taken 2211 non-allergenic chemical compounds with a filter of non-peptidic; No IgE; No histamine; No hypersensitivity; No allergy; No Cancer from the IEDB database. The chemical compounds with allergenic properties were considered as a positive dataset (allergens), and compounds with non-allergenic properties were taken as a negative dataset (non-allergens). Further, compound Ids were used to download the 2D and 3D structure files for 519 allergen and 2211 non-allergen chemical compounds. However, out of 2730 compounds, only 403 positive and 1074 negative compound structures were available in ChEBI. Final dataset contains 403 positive and 1074 negative chemical compounds. This dataset was divided into 80:20 ratio, where 80% of the data was used for training and 20% data validation. Our training dataset comprises of 320 allergens and 859 non-allergens, whereas our validation dataset comprises of 83 allergens and 215 non-allergens.

### Generation of Descriptors

The chemical descriptors/features of allergen and non-allergen chemical compounds were computed using PaDEL software.^**22**^ It can compute number of molecular descriptors, such as 2D, 3D and different types of fingerprints for a single chemical compound. It has computed 729 2D descriptors, 431 3D descriptors, and 16092 binary fingerprint-based (FP) descriptors for the 403 allergen and 1074 non-allergen chemical compounds. These 2D, 3D, and FP descriptor files were further used to develop different machine learning models.

### Dataset Preprocessing

These descriptors have values in different range, we perform preprocessing to normalize values of these descriptors. In this study, we used a well-established standard scaler method. The normalization and preprocessing were performed using a standard scaler package of scikit learn, i.e., sklearn.preprocessing.StandardScaler, which is based on a z-score normalization algorithm.^**25**^

### Selection of Descriptors

It has been shown in past studies, that all the descriptors are not significant.^**26, 27**^ Hence, it is important to find out the most relevant features from the vast number of descriptors. There are many feature selection techniques available; however, in this study, we have used the variance threshold-based method, correlation-based method and SVC-L1-based feature selection technique to select the significant features. Firstly, we have removed the low variance features from all descriptor files using the VarianceThreshold feature selection method from the sklearn package.^**25**^ It is used to filter-out low variance 2D, 3D and FP descriptors from the positive and negative data. Initially, there were 729 2D, 431 3D, and 16092 FP descriptors. After removing low variance features, we were left with 286 2D, 362 3D, and 1957 FP descriptors.

Secondly, we have used the correlation-based feature selection method for the removal of highly correlated features. We developed a python script to compute the pairwise correlation of all descriptors of each dataset. Then we have removed those features which were having a correlation of greater than or equal to 0.6. In this way, remaining were those features which have a correlation less than 0.6 with each other. As a result, we were left with 34 descriptors out of 286 descriptors for 2D, 8 descriptors out of 362 descriptors for 3D, and 210 descriptors out of 1957 FP descriptors. In order to get a highly significant feature set, we further tried to reduce the feature vector size using the most popular feature selection method, i.e., SVC-L1. ^**28, 29**^ It simultaneously performs a number of methods to select the best features from a large feature vector. It selects the non-zero coefficients and then implements the L1 penalty to choose the relevant features from the large feature vector to reduce dimensions. Based on this technique, we get the most important feature set, i.e., 14 descriptors out of 34 descriptors for 2D, 6 out of 8 descriptors for 3D and 22 FP descriptors out of 957 descriptors. The information of the regarding the selected descriptors is tabulated in Supplementary Table 1.

### Machine learning models

In this study, different machine learning techniques have been used for the classification of allergen and non-allergen chemical compounds. Logistic Regression (LR), k-nearest neighbors (KNNs), Decision Tree (DT), Gaussian Naive Bayes (GNB), XGBoost (XGB), Support Vector Classifier (SVC), and Random Forest (RF) were implemented to develop the classification models.

### Cross-validation and Evaluation Parameters

Several studies in the past have used 80:20 ratio for the division of the complete dataset.^**12, 29**^ In present study, to evaluate the developed machine learning models, we have applied 5-fold cross-validation on 80% of the training data for the internal training, testing and model evaluation. ^**30, 31**^ In 5-fold CV, the training data is divided into 5-sets, where four sets were used for the training and fifth set was utilized for the testing purposes. The same process is repeated five times, so that each set of positive and negative data is used for training and testing purposes. The performance of machine learning models was evaluated using the standard evaluation parameters. Threshold dependent and independent parameters both were used to measure the performance. Sensitivity (Sens), Specificity (Spec), Accuracy (Acc), Matthews correlation coefficient (MCC) are threshold dependent parameters, whereas the area under receiver operating characteristic curve (AUC) is a threshold independent parameter. These performance evaluation parameters are well-defined in the literature and have been extensively used in assessing the performance of the model.^**12, 32, 33**^

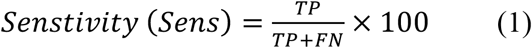

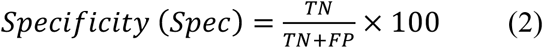

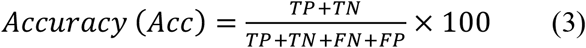

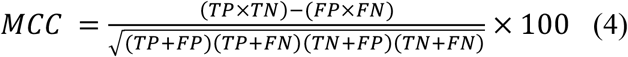

where FP, FN, TP, and TN are false positive, false negative, true positive, and true negative predictions, respectively.

## Results

### Fingerprints based analysis

In order to understand, importance of each fingerprint in classification of allergens and nonallergens, we compute prediction ability of each fingerprint. We used our in-house scripts, to check discrimination ability fingerprint (FP) based descriptors calculated by PaDEL. We ranked the fingerprints according to their probabilities for correctly classifying the chemical as allergen and non-allergen. Based on ranking, we identified most important 20 fingerprints. Ten fingerprints are highly present in allergens and were called positive fingerprints, where as other 10 which are highly present in non-allergens were called as negative fingerprints. Figure 2 depicts the frequency of top 10 positive and 10 negative fingerprints in allergens and nonallergens. These 10 positive fingerprints are highly abundant in allergens but negligible in nonallergens. Similarly, 10 negative fingerprints are highly abundant in non-allergens but negligible in allergens. The complete information regarding these top fingerprints is provided in Supplementary Table 2.

**Figure 2:**
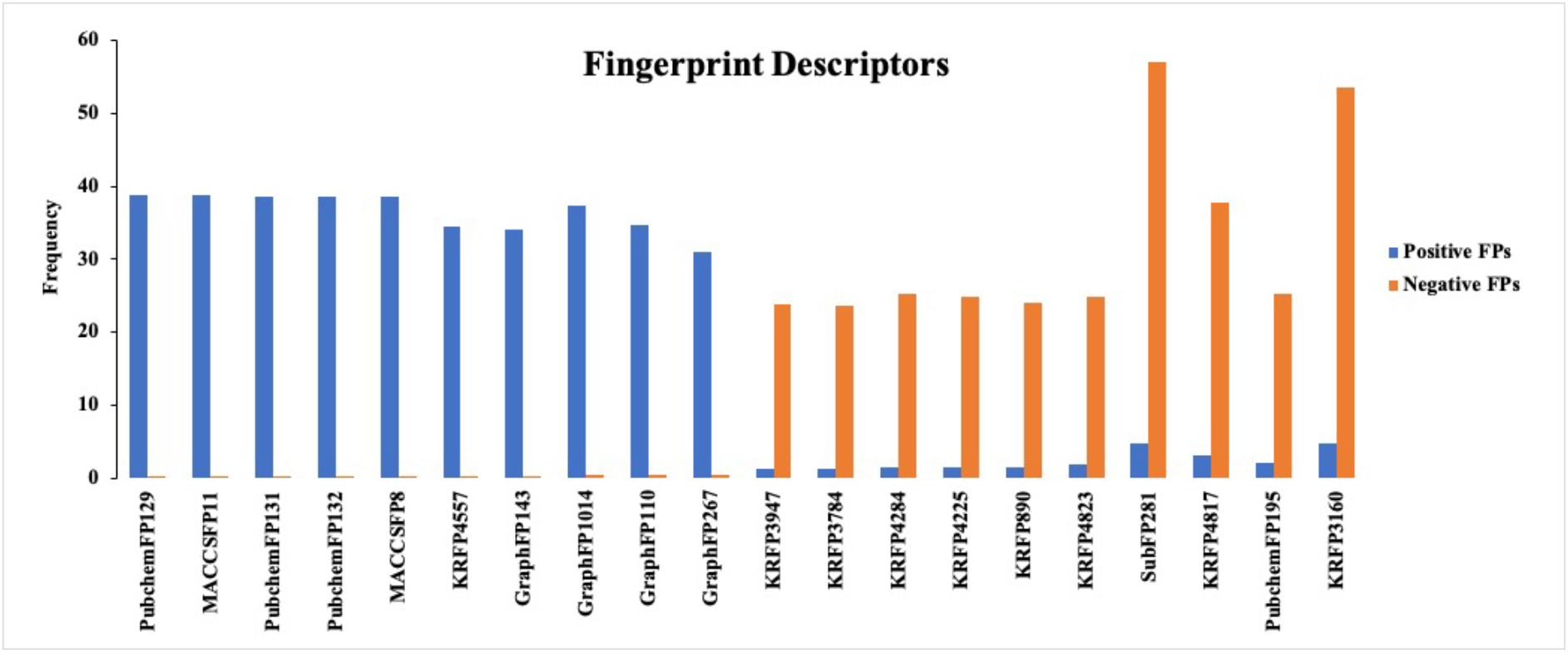
Shows frequency of top 10 positive and 10 negative fingerprints in allergens and non-allergens.

### Prediction Models using 2D/3D/FP Descriptors

Several models have been developed for predicting chemical allergens using different kinds of chemical descriptors like 2D, 3D and FP descriptors. Several machine learning approaches have been used for developing prediction models, it includes RF, KNN, XGB, SVC, LR, GNB, and DT. The models developed by using these machine learning techniques were optimized by tuning different parameters on training dataset using five-fold cross validation. Firstly, we have developed the machine learning models using 14 descriptors selected from 2D descriptors. The model based on the XGB algorithm performed better than other classifiers and achieved maximum AUC 0.90 and 0.89 on the training and validation datasets, respectively. Similarly models were developed using 6 features selected from 3D descriptors. Random forest (RF) based model outperformed the other methods and achieved maximum AUC 0.88 and 0.85 on the training and validation datasets, respectively (Table 1). In order to develop models using fingerprint, we selected 22 out of total 16092 fingerprints. Our RF-based model on 22 fingerprints achieved maximum AUC 0.92 and 0.92 on the training and validation datasets, respectively (Table 2).

**Table 1:** The performance of ML-based models developed using 14 (2D) descriptors and 6 (3D) descriptors.

| 2D Descriptors |  |  |  |  |  |  |  |  |  |  |
| --- | --- | --- | --- | --- | --- | --- | --- | --- | --- | --- |
| ML | Training |  |  |  |  | Validation |  |  |  |  |
|  | Sens | Spec | Acc | AUC | MCC | Sens | Spec | Acc | AUC | MCC |
| **XGB** | 81.68 | 82.11 | 81.99 | 0.90 | 0.60 | 80.25 | 76.64 | 77.63 | 0.89 | 0.52 |
| **KNN** | 81.37 | 81.99 | 81.82 | 0.90 | 0.59 | 81.48 | 79.91 | 80.34 | 0.88 | 0.57 |
| **RF** | 81.99 | 81.29 | 81.48 | 0.90 | 0.59 | 83.95 | 81.31 | 82.03 | 0.90 | 0.61 |
| **LR** | 80.12 | 81.05 | 80.80 | 0.88 | 0.57 | 81.48 | 77.57 | 78.64 | 0.88 | 0.54 |
| **DT** | 79.50 | 79.65 | 79.61 | 0.85 | 0.55 | 67.90 | 76.64 | 74.24 | 0.80 | 0.42 |
| **GNB** | 78.57 | 78.36 | 78.42 | 0.86 | 0.53 | 81.48 | 77.57 | 78.64 | 0.87 | 0.54 |
| **SVC** | 78.26 | 77.78 | 77.91 | 0.87 | 0.52 | 85.19 | 78.04 | 80.00 | 0.88 | 0.58 |
| 3D Descriptors |  |  |  |  |  |  |  |  |  |  |
| **RF** | 79.14 | 78.69 | 78.81 | 0.88 | 0.54 | 75.33 | 81.19 | 79.66 | 0.85 | 0.53 |
| **KNN** | 77.61 | 77.87 | 77.80 | 0.85 | 0.51 | 68.83 | 80.73 | 77.63 | 0.83 | 0.47 |
| **XGB** | 76.69 | 76.11 | 76.27 | 0.86 | 0.49 | 77.92 | 79.36 | 78.98 | 0.86 | 0.53 |
| **SVC** | 73.31 | 72.25 | 72.54 | 0.81 | 0.42 | 62.34 | 73.85 | 70.85 | 0.77 | 0.33 |
| **LR** | 68.41 | 71.31 | 70.51 | 0.73 | 0.36 | 70.13 | 72.48 | 71.86 | 0.76 | 0.38 |
| **GNB** | 68.41 | 70.61 | 70.00 | 0.75 | 0.36 | 64.94 | 73.39 | 71.19 | 0.75 | 0.35 |
| **DT** | 69.33 | 68.38 | 68.64 | 0.76 | 0.34 | 71.43 | 60.55 | 63.39 | 0.72 | 0.28 |

**Table 2:** The performance of ML-based models developed using 22 (FP) descriptors.

| ML | Training |  |  |  |  | Validation |  |  |  |  |
| --- | --- | --- | --- | --- | --- | --- | --- | --- | --- | --- |
|  | Sens | Spec | Acc | AUC | MCC | Sens | Spec | Acc | AUC | MCC |
| **RF** | 85.06 | 85.11 | 85.10 | 0.92 | 0.66 | 86.67 | 85.52 | 85.81 | 0.92 | 0.67 |
| **XGB** | 85.37 | 85.11 | 85.18 | 0.92 | 0.66 | 85.33 | 85.52 | 85.47 | 0.90 | 0.66 |
| **LR** | 83.84 | 83.82 | 83.83 | 0.91 | 0.64 | 81.33 | 81.45 | 81.42 | 0.86 | 0.58 |
| **SVC** | 83.54 | 83.00 | 83.15 | 0.91 | 0.62 | 82.67 | 80.54 | 81.08 | 0.86 | 0.58 |
| **KNN** | 82.93 | 83.12 | 83.07 | 0.90 | 0.62 | 85.33 | 80.54 | 81.76 | 0.87 | 0.60 |
| **GNB** | 79.57 | 79.37 | 79.42 | 0.88 | 0.55 | 70.67 | 81.45 | 78.72 | 0.83 | 0.49 |
| **DT** | 79.88 | 78.90 | 79.17 | 0.86 | 0.54 | 77.33 | 76.92 | 77.03 | 0.83 | 0.49 |

### Prediction Models using Hybrid Features

In addition to developing models using each type of descriptors, we also developed models using selected features of all types of descriptors. Our hybrid model developed using 42 features that contain all three types of descriptors, i.e., 2D (14 features), 3D (6 features) and FP (22 features). Our RF-based model had achieved maximum AUC 0.94 and 0.93 on the training and validation datasets, respectively with balanced sensitivity and specificity. We have also developed the model using only 2D and FP descriptors to check the performance of the model. As there were only 6 (3D) descriptors, we have excluded them and have developed the model with only 36 features (14 (2D) and 22 (FP) descriptors). As depicted in Table 3, the RF-based model has obtained an AUC of 0.94 on the training dataset and 0.93 on the validation dataset. It indicates that 36 features are sufficient to achieve highest performance, which is best model.

**Table 3:** The performance of ML-based hybrid models developed after combining all descriptors.

| **42 Hybrid Descriptors (2D+3D+FP)** |  |  |  |  |  |  |  |  |  |  |
| --- | --- | --- | --- | --- | --- | --- | --- | --- | --- | --- |
| **ML** | **Training** |  |  |  |  | **Validation** |  |  |  |  |
|  | **Sens** | **Spec** | **Acc** | **AUC** | **MCC** | **Sens** | **Spec** | **Acc** | **AUC** | **MCC** |
| **RF** | 85.63 | 86.00 | 85.90 | 0.94 | 0.68 | 87.95 | 82.55 | 84.07 | 0.93 | 0.66 |
| **SVC** | 84.06 | 84.01 | 84.03 | 0.91 | 0.64 | 93.98 | 78.77 | 83.05 | 0.92 | 0.66 |
| **KNN** | 84.06 | 83.66 | 83.77 | 0.92 | 0.63 | 80.72 | 81.60 | 81.36 | 0.92 | 0.58 |
| **XGB** | 83.75 | 83.78 | 83.77 | 0.92 | 0.63 | 85.54 | 79.72 | 81.36 | 0.92 | 0.60 |
| **LR** | 83.75 | 83.08 | 83.26 | 0.91 | 0.62 | 85.54 | 82.08 | 83.05 | 0.89 | 0.63 |
| **GNB** | 84.06 | 79.70 | 80.88 | 0.89 | 0.59 | 73.49 | 83.02 | 80.34 | 0.87 | 0.54 |
| **DT** | 79.06 | 78.76 | 78.85 | 0.87 | 0.53 | 81.93 | 80.66 | 81.02 | 0.88 | 0.58 |
| **36 Hybrid Descriptors (2D+FP)** |  |  |  |  |  |  |  |  |  |  |
| **RF** | 87.5 | 87.28 | 87.34 | 0.94 | 0.71 | 84.34 | 83.02 | 83.39 | 0.93 | 0.63 |
| **XGB** | 85 | 84.95 | 84.96 | 0.93 | 0.66 | 81.93 | 81.13 | 81.36 | 0.91 | 0.59 |
| **LR** | 84.06 | 84.48 | 84.37 | 0.91 | 0.64 | 84.34 | 83.96 | 84.07 | 0.90 | 0.64 |
| **KNN** | 83.44 | 84.13 | 83.94 | 0.93 | 0.63 | 81.93 | 82.08 | 82.03 | 0.91 | 0.60 |
| **SVC** | 84.06 | 83.08 | 83.35 | 0.90 | 0.63 | 86.75 | 81.60 | 83.05 | 0.91 | 0.63 |
| **GNB** | 84.06 | 79.00 | 80.37 | 0.88 | 0.58 | 77.11 | 83.96 | 82.03 | 0.88 | 0.58 |
| **DT** | 80 | 79.00 | 79.27 | 0.86 | 0.54 | 86.75 | 76.89 | 79.66 | 0.88 | 0.58 |

### Webserver interface

We have developed a user-friendly web server named ChAlPred for the prediction of chemicals as allergens and non-allergens. In this server, we have provided the three modules: (i) predict, (ii) draw and (iii) analog design module. The Predict module allows the user to submit the chemical compounds in different formats, such as SMILE, SDF and MOL formats, to predict whether the chemical could be allergenic or non-allergenic. The Draw module allows the user to draw or modify a molecule in an interactive way using Ketcher^**34**^ and submit the molecule to the machine learning models to predict whether the modified compounds will be allergenic or not. The Analog design module can be used to generate analogs based upon a combination of a given scaffold, building blocks and linkers. The server subsequently predicts the generated analogs as allergenic or non-allergenic. The web server has been designed using a responsive HTML template and browser compatibility for different OS systems.

### Case Study: Potential allergenic FDA-approved drugs

In order to identify the FDA-approved drugs that can cause allergic reactions to the person, we have downloaded the total of 2675 FDA drug molecules from the DrugBank Database.^**35**^ Out of 2675, we have only considered 1102 drugs which are approved. From 1102 drug molecules, the 2D structures were available only for 842 drugs. Finally, we have the structures of 842 FDA-approved drug molecules, which were used to identify that which drug molecules could be allergenic and non-allergenic. We have used the RF-based machine learning model of the Predict module on the “ChAlPred” web server. The prediction was made using the default parameters. The model has predicted 114 drug molecules to be allergenic. Several studies done in the past have also supported our findings that some of these drugs can cause allergy in the patient when administered. We have identified 20 drug molecules which are used to cure some diseases but also tend to cause allergic symptoms. Table 4 depicts the information of the drug molecules which cause some allergic reactions.

**Table 4:** FDA-approved drug molecules predicted by our server (ChAlPred) causing allergic symptoms.

| Drug Bank ID | FDA-Approved Drugs | Prediction | Allergic Symptoms |
| --- | --- | --- | --- |
| DB01112 | Cefuroxime | Allergen | Anaphylactic Reaction <sup>36</sup> |
| DB00421 | Spironolactone | Allergen | Skin allergy , drug rash with eosinophilia and systemic symptoms (DRESS) induced by spironolactone <sup>37</sup> |
| DB00859 | Penicillamine | Allergen | Skin allergy <sup>38</sup> |
| DB05013 | Ingenol mebutate | Allergen | Skin allergy <sup>39</sup> |
| DB01007 | Tioconazole | Allergen | Contact hypersensitivity <sup>40</sup> |
| DB06209 | Prasugrel | Allergen | Hypersensitivity Skin Reaction <sup>41</sup> |
| DB01330 | Cefotetan | Allergen | Cefotetan-induced anaphylaxis <sup>42</sup> |
| DB01331 | Cefoxitin | Allergen | Allergic reactions <sup>43</sup> |
| DB04854 | Febuxostat | Allergen | Hypersensitivity Reactions (HSRs) <sup>44</sup> |
| DB09212 | Loxoprofen | Allergen | Type I allergic reaction, eosinophilic coronary periarthritis <sup>45</sup> |
| DB00973 | Ezetimibe | Allergen | Angioedema allergic reaction <sup>46</sup> |
| DB00390 | Digoxin | Allergen | Cutaneous hypersensitivity <sup>47</sup> |
| DB00493 | Cefotaxime | Allergen | Immediate Hypersensitivity Reactions <sup>48</sup> |
| DB01150 | Cefprozil | Allergen | Immediate Hypersensitivity Reactions <sup>48</sup> |
| DB00833 | Cefaclor | Allergen | Immediate Hypersensitivity Reactions <sup>48</sup> |
| DB00689 | Cephaloglycin | Allergen | Immediate Hypersensitivity Reactions <sup>48</sup> |
| DB00438 | Ceftazidime | Allergen | Immediate Hypersensitivity Reactions <sup>48</sup> |
| DB00267 | Cefmenoxime | Allergen | Immediate Hypersensitivity Reactions <sup>48</sup> |
| DB00703 | Methazolamide | Allergen | Skin allergy <sup>49</sup> |
| DB13154 | Parachlorophenol | Allergen | Allergic contact dermatitis <sup>50</sup> |

## Discussion and Conclusion

One of the major challenges in the field of drug discovery is side effect or adverse reaction of drugs. In the past, number of drugs have been already withdrawn from market due to their adverse effects. A wide range of toxicities are responsible for side-effect of drugs, it may be cytotoxicity, immuno-toxicity, hemo-toxicity, liver toxicity or allergenicity.^**51**^ Identification of toxicity is costly, time consuming and tedious task. Thus, there is a need to predict these toxicities using in silico methods. In the past, various tools have been developed to estimate the toxicity of the chemicals such as The Toxicity Estimation Software Tool (TEST)^**52**^ VegaQSAR, and Toxtree^**53**^. These tools are based on the Quantitative Structure-Activity Relationships (QSARs) model for toxicity prediction of the chemical molecules. Machine learning-based tools such as ToxiM, developed by Sharma et al., predict the toxicity and toxicity-related properties of small chemical molecules using machine learning approaches^**19**^, ProTox-II.^**54**^

In contrast, no tool have been developed for predicting allergenicity of the chemical molecules. In this current work, we have collected chemical compounds with their well-defined molecular descriptors utilising publicly available databases such as IEDB and ChEBI. The data yielded several descriptors which was reduced using various feature selection methods. We sorted the most important feature set, i.e., 14 descriptors for 2D, 6 descriptors for 3D and 22 FP descriptors. Based on these selected features (14 2D and 22 FP) we have successfully employed several machine learning approach and found that RF attained a maximum AUC of 0.94 and 0.93 on training as well as validation dataset. In addition, fingerprints based analysis suggests that two positive fingerprints, i.e., PubChemFP129 (Extended Smallest Set of Smallest Rings (ESSSR) ring set >= 1 any ring size 4) and GraphFP1014 are highly present in allergenic chemical compounds, and three negative fingerprint, i.e., Klekota-Roth fingerprints (KRFP890 ([!#1][NH]C(=O)[CH3], KRFP3160 (C1CCOCC1)) and Substructure fingerprint (SubFP281 [OX2;$([r5]1@C@C@C(O)@C1),$([r6]1@C@C@C(O)@C(O)@C1)]) are abundant in non-allergenic chemical compounds. FDA-approved drugs analysis have shown that few drugs which are used for treatment of certain diseases, are also causing allergy as the side effect. Literature evidences have shown that administration of FDA-approved drugs such as, Cefuroxime^**36**^, Spironolactone ^**37**^, Penicillamine^**38**^ can cause allergic reactions like, skin allergies, anaphylactic reactions, hypersensitivity. For instance, a case report has shown that 60 year old patient was experiencing anaphylactic reaction after given an antibiotic cefuroxime.^**55**^ Another report by Kinsara has shown that Spironolactone, a potassium sparing diuretic was given to a patient which was diagnosed with idiopathic cardiomyopathy, and he developed a macular rashes on both the arms (https://asclepiusopen.com/journal-of-clinical-cardiology-and-diagnostics/volume-1-issue-2/1.php). A clinical study by Zhu et al, reported that the patients with Wilson disease were given D-penicillamine (DPA) medication at first, but later they developed neurological symptoms as well as allergies.^**38**^

We can see that these medications can cause a variety of allergic reactions in patients, some of which can be fatal. To prevent these problems, there is a need for predicting the allergenicity of chemical compounds before using them for treatment purposes. Eventually, we built a freely available webserver namely ChAlPred, for predicting allergenic and non-allergenic chemical compounds using machine learning techniques based on their 2D, 3D and FP molecular descriptors. We hope that this study will be helpful in the future for designing the drug molecules with no allergenic properties.

## Abbreviations

IEDB: The Immune Epitope Database
ChEBI: Chemical Entities of Biological Interest
FDA: The Food and Drug Administration
HTML: The HyperText Markup Language
SMILES: The simplified molecular-input line-entry system

## Acknowledgement

The authors are thankful to Department of Science and Technology (DST-INSPIRE), Department of Biotechnology (DBT) for fellowships, financial support, and IIIT-Delhi for providing the infrastructure and facilities.

## Authors Contribution

**Conception and design:** Neelam Sharma, Gajendra P. S. Raghava.

**Development of methodology:** Neelam Sharma, Sumeet Patiyal, Anjali Dhall, Gajendra P. S. Raghava.

**Acquisition of data:** Neelam Sharma.

**Analysis and interpretation of data and results:** Neelam Sharma, Sumeet Patiyal, Anjali Dhall, Gajendra P. S. Raghava.

**Webserver Implementation:** Neelam Sharma, Sumeet Patiyal.

**Writing, reviewing, and revision of the manuscript:** Neelam Sharma, Anjali Dhall, Naorem Leimarembi Devi, Gajendra P. S. Raghava.

## Data Availability

All the datasets generated for this study are available at the “ChAlPred” webserver, https://webs.iiitd.edu.in/raghava/chalpred/dataset.php.

